# Genetic and environmental contributions to ReHo and fALFF in early adolescence vary across brain regions

**DOI:** 10.1101/2025.01.31.635399

**Authors:** Lachlan T Strike, Katie L McMahon, Sarah E Medland, Greig I de Zubicaray

## Abstract

Research on genetic and environmental influences on brain function generally focuses on connections between brain areas. A different yet unexplored approach is to examine activity within local brain regions. We investigated the influence of genes and environmental effects on two specific measures of local brain function: Regional Homogeneity (ReHo) and fractional Amplitude of Low-Frequency Fluctuations (fALFF). Participants were drawn from a sample of adolescent twins on two occasions (mean ages 11.5 and 13.2 years, *N* = 278 and 248). Results showed that genetic and environmental factors influenced brain function in almost all 210 cortical regions examined. Moreover, genetic and common environmental factors influencing ReHo and fALFF values at wave 1 (9-14 years) also influenced values at wave 2 (10-16 years) for many regions. However, the influence of genetic and common environmental factors varied across the cortex, exhibiting different patterns in different regions. Furthermore, we found new (i.e., independent) genetic and environmental influences on brain activity at wave 2, again with regional patterns. Exploratory analyses found weak associations between anxiety and depressive symptoms and local brain function in several regions of the temporal lobe. These findings are consistent with similar studies of other resting-state functional MRI metrics (i.e., functional connectivity).

---

Characterising genetic and environmental influences on adolescent brain function is vital to understanding this often turbulent developmental period. In recent years, resting-state functional magnetic resonance imaging (rs-fMRI) has emerged as a powerful tool for investigating intrinsic brain function. Two key metrics derived from rs-fMRI – fractional Amplitude of Low-Frequency Fluctuations (fALFF) and Regional Homogeneity (ReHo) - provide insights into local neural dynamics (Zang et al. 2004; Zou et al. 2008; Lv et al. 2018). fALFF reflects the strength of low-frequency oscillations (i.e., local activity), while ReHo measures the temporal coherence of neighbouring voxels within specific brain regions (i.e., local connectivity).

ReHo and fALFF have been widely adopted as potential predictors of attention deficit hyperactivity disorder (Zhu et al. 2008; Alonso et al. 2014; Tan et al. 2017) and have been examined as neurobiological signatures of anxiety and depression (Oathes et al. 2015; Shen et al. 2020; Zhao et al. 2023). In addition, ReHo and fALFF measures have shown alterations in young patients with anorexia nervosa (Seidel et al. 2019; Seidel et al. 2024) and adolescents involved in contact sports (Li et al. 2022; Zuidema et al. 2024). Moreover, variations in ReHo and fALFF brain function measures have been related to cognitive ability in children and adults (Yang et al. 2015; Koyama et al. 2020; Kearney et al. 2023) as well as Alzheimer’s disease and mild cognitive impairment groups (Cha et al. 2015).

Genetic and environmental factors likely underlie variation in ReHo and fALFF measures of brain function, but work to date is limited (Adhikari et al. 2020). Studies examining genetic influences on rs-fMRI measures of brain function have typically focused on functional connectivity (i.e., the temporal coherence of voxels in distant brain regions), a related but separate rs-fMRI metric to ReHo and fALFF. Studies have shown a wide range of heritability estimates (i.e., the proportion of phenotypic variance attributed to genetic effects) for functional connectivity in adults (up to ≈80%). However, estimates are typically small to moderate (Glahn et al. 2010; Sinclair et al. 2015; Yang et al. 2016; Adhikari et al. 2018; Barber et al. 2021). Heritability estimates for rs-fMRI in children and adolescents are limited (van den Heuvel et al. 2013; Fu et al. 2015; Achterberg et al. 2018; Teeuw, Brouwer, Guimaraes, et al. 2019) and again show a wide range of estimates (up to ≈80%). Interestingly, studies in adults and children have demonstrated regional variation in the level of genetic influence across cortical networks, though these patterns show little consistency between studies. This inconsistency across studies is likely due to the relatively low reproducibility of rs-fMRI connectivity measures (Golestani et al. 2017; Cahart et al. 2023), as reproducibility typically places an upper limit on heritability estimates.

A further unanswered question of functional brain development is whether the genetic and environmental factors influencing brain function change over time. In a sample of adolescent twins scanned at ages 13 and 18, Teeuw et al. (2019) reported that genetic influences on functional connectivity were generally stable between ages 13 and 18, with little evidence of fluctuating genetic influences. However, as the authors note, the reduced sample size at age 13 compared to age 18 (due to a greater incidence of dental braces and head motion) limited the author’s ability to examine changes in genetic and non-genetic influence across time. Studies of structural brain phenotypes have suggested the presence of independent genetic factors during childhood and adolescence (van Soelen et al. 2012; Teeuw, Brouwer, Koenis, et al. 2019), highlighting the need to examine functional brain measures further.

Though related to functional connectivity, ReHo and fALFF inform on different aspects of neural activity, and it cannot be assumed that they share a similar magnitude of genetic and environmental influences as functional connectivity estimates. In addition, they demonstrate greater reproducibility than functional connectivity measures (Golestani et al. 2017; Cahart et al. 2023). As such, we estimate the heritability of ReHo and fALFF values in a dataset of young adolescent twins scanned over two waves (mean age 11.4 and 13.1 years, respectively) while carefully examining other variance sources (i.e., shared environment and unique environment). We also investigate whether the genetic (and non-genetic) factors influencing ReHo and fALFF continue between the two waves. We expect to find patterns of genetic influence similar to those reported by Teeuw et al. (2019) for functional connectivity measures, specifically higher heritability estimates for ReHo and fALFF values in regions associated with visual, frontoparietal and salience networks. Further, we expect these genetic influences will overlap between waves 1 and 2, but that small to moderate novel genetic influences will exist in wave 2. As an exploratory analysis, we examine behavioural correlates of ReHo and fALFF values, aiming to replicate past findings with mood and cognitive phenotypes (Oathes et al. 2015; Wang et al. 2016; Shen et al. 2020).

## Materials and methods

### Participants

Participants were from the Queensland Twin Adolescent Brain (QTAB) longitudinal brain development study. The QTAB study (O’Callaghan et al. 2021; Hansell et al. 2022; Strike et al. 2023) focused on late childhood/early adolescence, with brain imaging, cognition, mental health, and early life/family demographics data collected over two waves (wave 1: 9-14 years, wave 2: 10-16 years). Co-twins attended the study centre together and were assessed in parallel, with one twin undergoing brain imaging while the other completed assessments outside of the scanner. Baseline measures (i.e., wave 1 data) were collected in 211 families (422 individuals, ages 9-14 years). One hundred fifty-two families returned for a second wave of data collection approximately 20 months later (304 individuals, ages 10-16 years). Fifty-nine families did not return for wave 2 (39 were no longer eligible [i.e., braces, moved interstate or poor T1w scans at wave 1] and 20 declined). The dataset examined here comprised 278 and 248 individuals at waves 1 and 2, respectively (196 individuals with ReHo/fALFF measures at both waves; see **Table 1** for full details). Unpaired twins in the final dataset (i.e., participants whose co-twin was excluded from analysis) were retained to improve estimates of means and variances. Zygosity in same-sex twins was determined by genotypic data (93% of twin pairs) or parental questionnaire (7% of twin pairs). The study was approved by the Human Research Ethics Committees at The University of Queensland and Children’s Health Queensland. Written consent was obtained from all participants and a parent/guardian. The QTAB study was a research project and did not include neuroradiological examinations.

**Table 1:** Demographics for the final QTAB dataset (after participant exclusions) and mean (standard deviation) values for mental health and cognition measures.

|  | Wave 1 | Wave 2 | Waves 1 & 2* |
| --- | --- | --- | --- |
| N (individuals) | 278 | 248 | 196 |
| Age (years) | 11.5 ± 1.33 | 13.18 ± 1.5 | W1: 11.54 ± 1.32<br>W2: 13.26 ± 1.5 |
| Sex (% female/male) | 54/46 | 55/45 | 58/42 |
| Pubertal Status (PDS score) | 1.88 | 2.47 | W1: 1.90<br>W2: 2.52 |
| Handedness (% right/left) | 83/17 | 86/14 | 87/13 |
| Zygoty (MZ/DZ complete twin pairs) | 56/49 | 56/51 | 37/36 |
| Mean framewise displacement (mm) | .19 | .18 | W1: .19<br>W2: .18 |
| SMFQ score | 4 (3) | 5 (4) | W1: 4 (3)<br>W2: 5 (4) |
| SPHERE-21 score | 9 (6) | 10 (6) | W1: 9 (6)<br>W2: 10 (6) |
| SCAS score | 24 (13) | 24 (15) | W1: 24 (13)<br>W2: 25 (16) |
| NIHTB-CB Crystallized Composite score | 91 (7) | 94 (6) | W1: 91 (6)<br>W2: 95 (6) |

Wave 1: 144 participants (34%) were excluded from the analysis due to excessive head motion in rs-fMRI (n = 119; based on mean framewise displacement > 0.3mm) or anatomical MRI scans (n = 10; based on visual inspection of motion severity), incomplete scan acquisition (n = 10,) or claustrophobia/declined participation (n = 5)).
Wave 2: 56 participants (18%) were excluded from the analysis due to excessive head motion in rs-fMRI (n = 54) or incomplete scan acquisition (n = 2).
\* Overlapping participants between waves 1 and 2 (after participant exclusions).
DZ dizygotic; MZ monozygotic; NIHTB-CB NIH Toolbox Cognition Battery (Weintraub et al. 2013; Heaton et al. 2014); SCAS Spence Children's Anxiety Scale (Spence et al. 2003); SMFQ Short Moods and Feelings (Angold et al. 1995); SPHERE-21 Somatic and Psychological Health Report (Couvry-Duchesne et al. 2017); PDS Puberty Development Scale (Petersen et al. 1988).

### Image acquisition and processing

We acquired anatomical and functional images on a 3T Siemens Magnetom Prisma paired with a 64-channel head coil at the Centre for Advanced Imaging, The University of Queensland. T1- weighted anatomical scans were acquired using a prototype MP2RAGE sequence [(Marques et al. 2010); voxel size = 0.8 mm isotropic, TR/TE/TI1/TI2 = 4000 ms/2.9 ms/700 ms/2220 ms, TA: 6:04; amplified background noise removed using an extension (https://github.com/benoitberanger/mp2rage) of Statistical Parametric Mapping 12 (SPM12; https://www.fil.ion.ucl.ac.uk/spm/software/spm12). Resting-state data (voxel size 2 mm isotropic, TR = 930 ms, TE = 30 ms, slices = 72, multiband acceleration factor = 6) were acquired with 327 volumes (5:13 min:sec) per run. Two runs with reversed phase-encoding directions (anterior-posterior [AP] and posterior-anterior [PA]) were acquired consecutively. Participants viewed an abstract visual stimulus to improve resting-state compliance (Vanderwal et al. 2015). Acquisition parameters were identical for wave 2 scans. The entire imaging protocol lasted approximately 1 hour (Strike et al. 2023), with the resting state and anatomical scans acquired at the beginning of the session.

### rs-fMRI preprocessing and ReHo/fALFF estimation

The resting-state datasets were processed using the Data Processing and Analysis for (Resting-State) Brain Imaging toolbox (DPABI v8.1; (Yan et al. 2016), which is based on the Data Processing Assistant for Resting-State fMRI (DPARSF v5.4; (Yan and Zang 2010)) and SPM12. For each wave, the first five volumes were discarded from each run, and the AP and PA time series were realigned to the first image of the AP run, and a mean image was generated and used to coregister the realigned series to the T1-weighted image. The T1-weighted image was next segmented using a tissue probability map (TPM) matched to the mean age of the Wave 1 sample (11.5 years (Wilke et al. 2008)) included in the CAT12 toolbox (Gaser et al. 2024) and the DARTEL toolbox (Ashburner 2007) was employed to create a custom group template from the grey (GM) and white matter (WM) images. Slice timing correction of the realigned time series was performed using the multiband acquisition values, followed by nuisance signal regression using WM, cerebrospinal fluid (CSF) global signals and head motion parameters (Friston 24 model). The time series were next normalised to Montreal Neurological Institute (MNI) atlas space using the DARTEL flow-field transformations, linearly detrended, and motion scrubbing performed using the criteria from Power et al. (2012). Finally, temporal bandpass filtering (0.01-0.08 Hz) was performed before calculating ReHo but not for fALFF. ReHo was calculated using Kendall’s rank-based coefficient of concordance (KCC-ReHo; (Zang et al. 2004) that assesses the synchronisation between a given voxel’s signal time course and those of its 26 nearest neighbouring voxels. fALFF (Zou et al. 2008) was calculated as the ratio of the power spectrum of low-frequency signal fluctuations (0.01-0.08 Hz) to that of the entire frequency range (0-0.25 Hz). ReHo and fALFF maps were Z-standardized and smoothed with a 6 mm FWHM isotropic Gaussian kernel.

Regional measures (i.e., the average time series within a region-of-interest [ROI]) were extracted for 105 cortical ROIs in each hemisphere of the Human Brainnetome Atlas (Fan et al. 2016); **Fig. 1**). Measures were estimated separately for AP and PA scans and then averaged to aid reliability (Cao et al., 2023). Participants with mean framewise displacement (Power et al. 2012) greater than .3 mm per rs-fMRI acquisition were excluded from the analysis (**Table 1**).

**Fig. 1.**
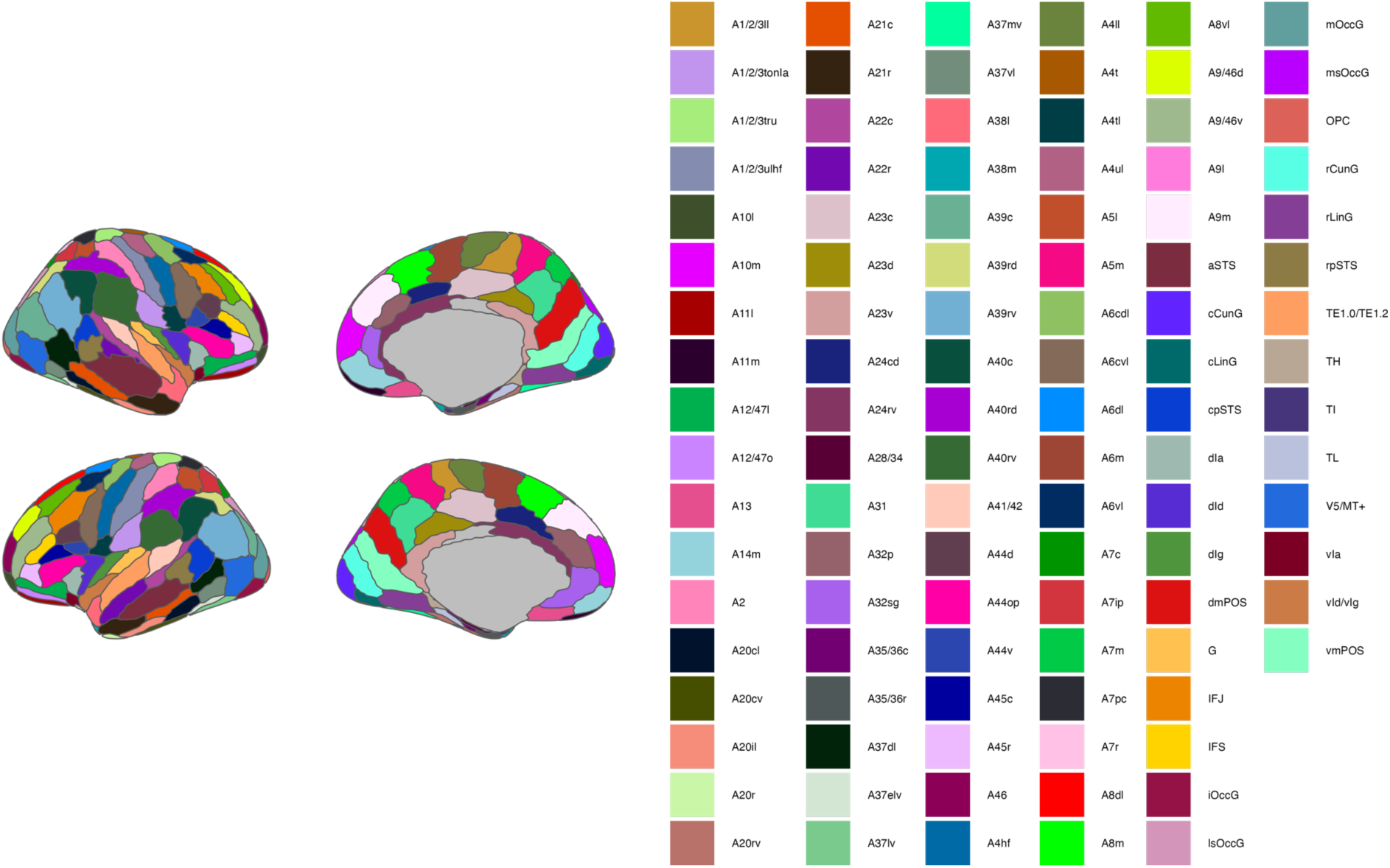
Cortical ROIs of the Brainnetome Atlas (105 ROIs per hemisphere). Note: The inferior temporal gyrus A20iv ROI is not presented (visible only through an inferior view of the brain).

### Mental health and cognition variables

A wide range of non-imaging measures were collected for QTAB participants (see Strike et al., (2023) for full details). Relevant to the current examination are symptoms of anxiety (Spence Children’s Anxiety Scale [SCAS; Spence et al., (2003)]), depression (Short Moods and Feelings Questionnaire [SMFQ; Angold et al.,(1995)], and anxiety/depression (Somatic and Psychological Health Report [SPHERE-21; Couvy-Duchesne et al., (2017)]). All anxiety and depressive symptom measures were assessed by participant self-report (total scores used for each scale). Cognitive ability was assessed using the NIH Toolbox Cognition Battery [NIHTB-CB](Weintraub et al. 2013; Heaton et al. 2014) administered via an iPad. The NIHTB-CB produces test-level and composite scores (i.e., Crystallized, Fluid, and Total Cognition Composite). We selected the Crystallized Cognition Composite as our cognitive variable of interest as it is the only composite score available for participants in both waves (the List Sorting Working Memory test was not collected at wave 2, preventing the creation of Fluid and Total Cognition Composite scores at wave 2).

### Saturated models

We began by fitting completely saturated bivariate models (which freely estimate all possible parameters) for ReHo or fALFF values of the 210 cortical regions of the Human Brainnetome Atlas (using wave 1 and 2 data). We then fit increasingly constrained saturated models to test twin modelling assumptions (i.e., equal means and variances across twin birth order and zygosity; see Evans et al., (1999)) and to check the dataset for errors. Saturated models were additionally used to examine covariate effects (age [years], sex assigned at birth [male = 0, female = 1], and mean framewise displacement [mm]). All covariates were carried forward for subsequent analyses regardless of effect size and significance. The significance of assumption tests and covariate effects was assessed through likelihood ratio tests comparing the fit between nested models (e.g., comparing models with and without a sex covariate (Grasby et al. 2017)). Correlations between monozygotic (MZ) and dizygotic (DZ) twin pairs and wave 1 and 2 ReHo/fALFF values were also estimated through saturated bivariate models.

### Bivariate models with Cholesky decompositions

We used the classical twin design using a path modelling approach (**Fig. 2a**; (Neale and Cardon 1992) to estimate genetic and environmental variance amounts in ReHo and fALFF ROI values. The classical twin design contrasts the observed covariance between MZ twins and DZ twins to partition the variance in a phenotype into four sources: additive genetic (A), non-additive genetic (D, e.g., dominance and epistasis), common or shared environment (C), and residual effects including idiosyncratic environmental factors and measurement error (E). However, as C and D are confounded in the classical twin design, they cannot be estimated simultaneously (i.e., only an ACE or ADE model is identifiable). The present study did not have sufficient power to discriminate between A and D effects (Keller et al. 2010), so models containing D effects were not considered.

**Fig. 2.**
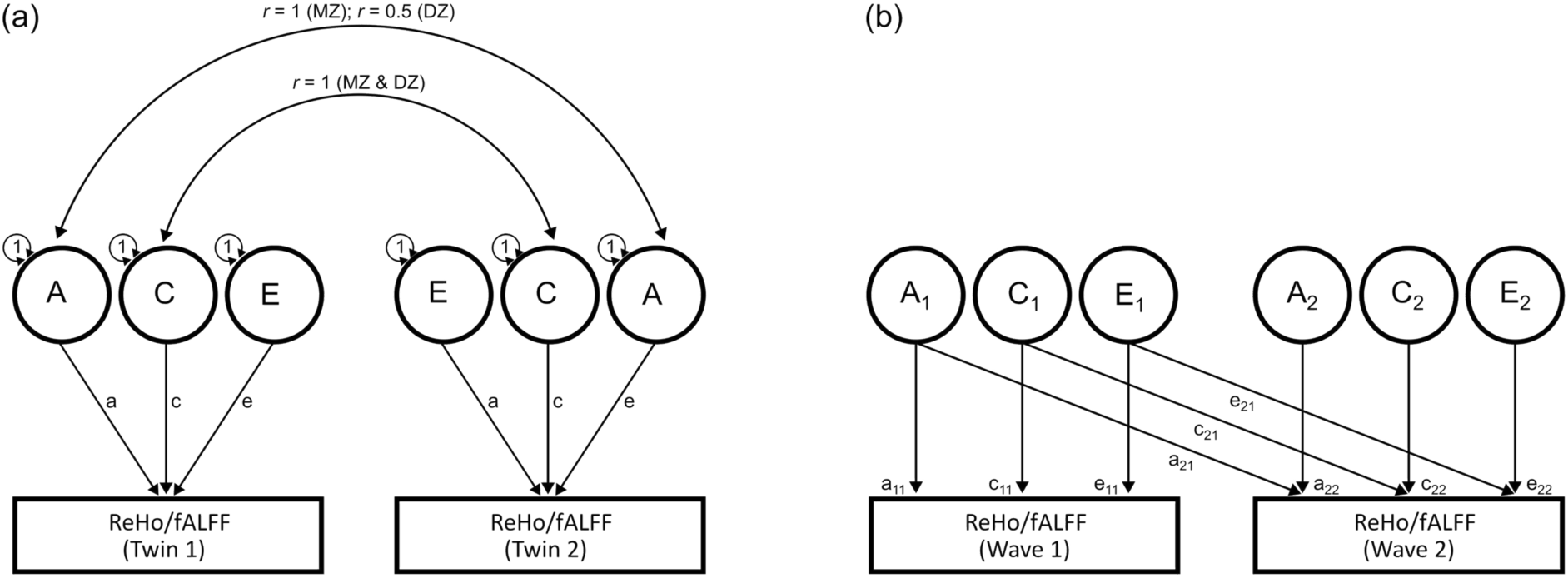
The classical twin design (**2a**) and bivariate extension specifying Cholesky decompositions (**2b;** shown for one twin only). In the classical twin design, correlations between additive genetic factors (A) are fixed at 1 for MZ and 0.5 for DZ twins, as MZ and DZ twins share 100% and (on average) 50% of their genetic material, respectively. For common environment effects (C), correlations are fixed at 1 for MZ and DZ twins. Unique environment effects (E) are uncorrelated between twin pairs, representing environmental influence affecting only one twin. Estimates of unique environment variance also include measurement error, as it is random and unrelated to twin similarity. The bivariate Cholesky design (**2b**) was used to decompose the (co)variance in wave 1 and 2 ReHo/fALFF measures into genetic and non-genetic sources. Latent factors (represented as circles) influence the observed variables (represented as rectangles) via paths (represented as arrows). The first latent genetic factor (*A*1) is assumed to explain the genetic variance in wave 1 ReHo/fALFF and can explain genetic variance in wave 2 ReHo/fALFF. The second genetic latent factor (*A*2) is uncorrelated with the first latent genetic factor and is assumed to explain the residual genetic variance in wave 2 ReHo/fALFF. Corresponding latent factors and decompositions are specified for common and unique environment variances. It is important to note that while the first latent factor (i.e., *A*1, *C*1, *E*1) can explain variance in multiple phenotypes, it cannot be interpreted as a common factor as it includes both common and specific variance (Loehlin 1996).

The multivariate extension of the classical twin design decomposes the covariance between two or more variables (or the same variable measured multiple times) into genetic and environmental components. Specifically, here, we used bivariate Cholesky decomposition analyses (**Fig. 2b**) to estimate genetic and environmental influences on variation in ReHo/fALFF values at wave 1 (9 – 14 years) and wave 2 (10 – 16 years). These analyses further examined the extent to which genetic and environmental effects continued from waves 1 to 2 or whether new effects emerged at wave 2. We tested the presence of new genetic influences at wave 2 by setting the wave 2 specific variance path (i.e., path *a*_22_ in **Fig. 2b**) to zero and assessing whether this significantly reduced model fit (via likelihood ratio test). In addition, we tested whether continuing genetic influences (i.e., influences overlapping between waves 1 and 2) diqered in strength between waves 1 and 2 by equating the paths originating from the first latent genetic factor (i.e., *a*_21_ = *a*_11_) and testing for a reduction in model fit. Similar tests were completed for common environmental eqects. Simplified sub-models containing AE, CE, and E variance sources were additionally fit to the data, and Akaike’s Information Criteria (AIC; Akaike (1974)) was used to compare the relative performance of these candidate models (models that minimise AIC are preferred).

### Associations with mental health and cognition measures

Exploratory analyses examined associations between ReHo and fALFF values and self-report measures of anxiety and depressive symptoms (SMFQ, SPHERE-21, SCAS) and cognition (Crystalized Composite Score) using wave 1 and 2 data. We fit linear mixed-effects models, specifying fixed effects of age, sex, mean framewise displacement, and wave (0 = wave 1, 1 = wave 2), with random effects specified for family (to control for relatedness) and participant (to control for repeat measurements [i.e., waves 1 and 2]).

### Software, multiple comparisons, and data availability

Saturated and bivariate twin models were fit using the maximum-likelihood structural equation modelling package OpenMx v2.21.8 (Neale et al. 2016; Boker et al. 2023) in R 4.4.1 (R Core Team 2024) using the NPSOL optimiser. Cortical plots were created using packages ggseg (Mowinckel and Vidal-Piñeiro 2020) and ggesgBrainnetome (Mowinckel 2021) in R 4.4.1 (R Core Team 2024). We used the Benjamini–Hochberg procedure to control the false discovery rate for the multiple comparisons (Benjamini and Hochberg 1995). The procedure was applied to significance tests of (1) covariate effects (i.e., age, sex, mean framewise displacement), (2) assumption testing (i.e., means and variance differences), (3) new genetic influences at wave 2 or (de)amplification of continuing genetic influences, and (4) exploratory analyses between ReHo/fALFF values and behavioural measures. MRI data for the QTAB project are freely available from the OpenNeuro data repository (Strike, Hansell, Miller, et al. 2022). The QTAB non-imaging phenotypes dataset (i.e., zygosity, mental health and cognition measures) requires a data transfer/usage agreement and is available from the Zenodo data repository (Strike, Hansell, Chuang, et al. 2022). Analysis code is available online at https://github.com/strikel/ReHo_fALFF_heritability.

## Results

### Preliminary analyses

Means, covariate effects, twin correlations, and phenotypic correlations between waves 1 and 2 for ReHo and fALFF values are presented in **STables 1-2**. Mean group-level ReHo and fALFF values (i.e., individual Z-scores averaged across all participants; also presented in **Fig. 3**) were lowest for parahippocampal gyrus and insular lobe regions and highest for occipital and parietal lobe regions. Assumption testing results showed a similar pattern of ReHo /fALFF values for co-twins and zygosity type. Age was positively associated with ReHo values of superior frontal gyrus ROIs, and negatively associated with ReHo values of temporal lobe ROIs. Associations between age and fALFF values were sparser and confined to the frontal and temporal lobes. Sex effects were minimal at wave 1; however, there was a marked increase in the number of ROIs showing effects of sex at wave 2, especially for fALFF (ReHo: 2 ROIs wave 1, 13 ROIs wave 2; fALFF: 3 ROIs wave 1, 54 ROIs wave 2). fALFF values were greater in females for ROIs in the frontal lobe and cingulate gyrus and greater in males for ROIs in the parietal/temporal lobes and insular gyrus. Mean framewise displacement was associated with ReHo or fALFF values for select ROIs across the whole cortex (ReHo: 15 ROIs wave 1, 37 ROIs wave 2; fALFF: 42 ROIs wave 1, 23 ROIs wave 2); however, associations were weak (up to a .08 change in ReHo or fALFF values associated with a 1 SD increase in mean FD).

**Fig. 3.**
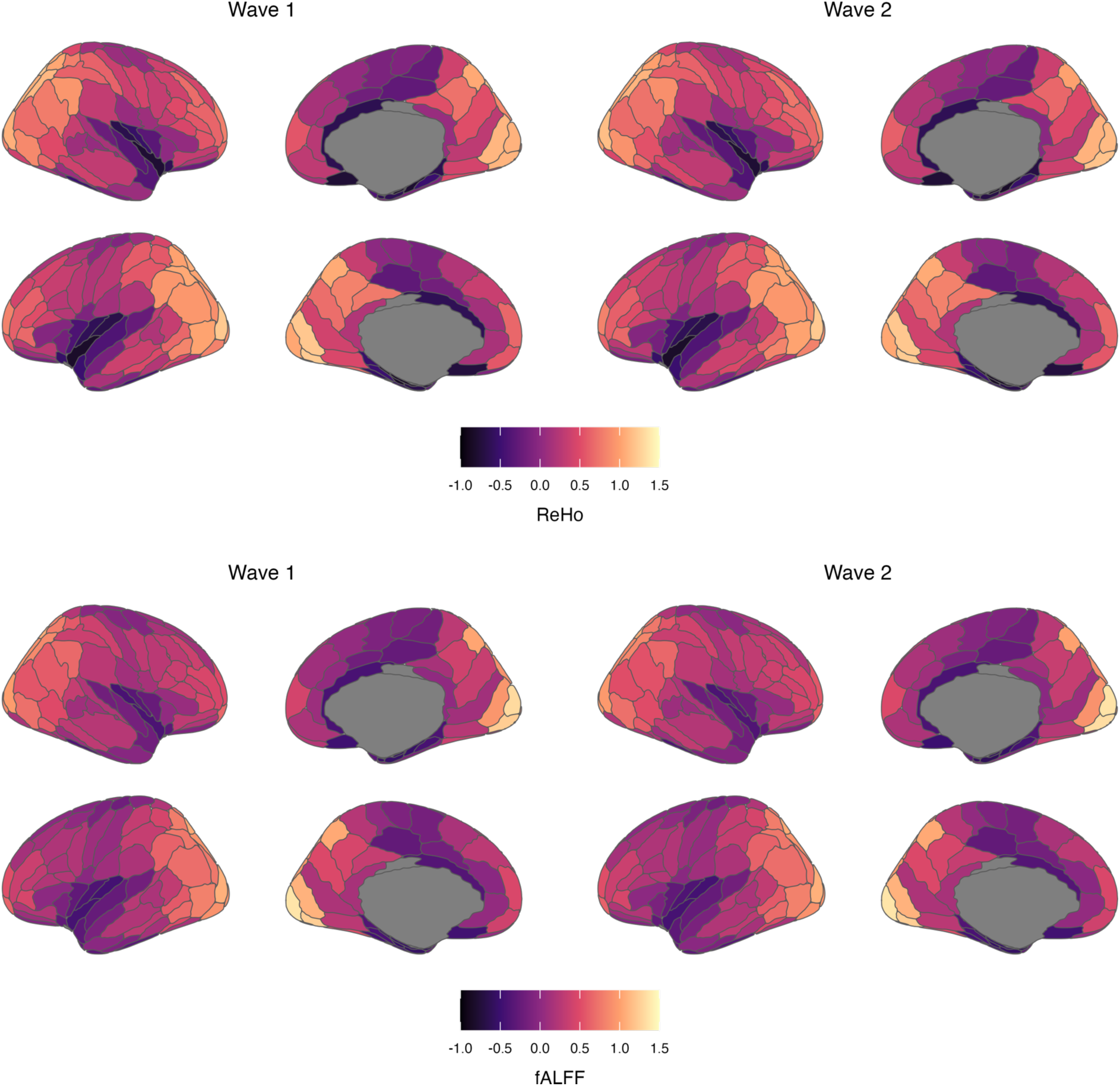
Group level ReHo and fALFF values for the QTAB dataset at waves 1 (*n* = 278 individuals) and 2 (*n* = 248 individuals). Values represent individual Z-scores from each participant, averaged across all participants at each wave).

Correlations between ReHo/fALFF values of twin pairs showed evidence for additive genetic influences (i.e., the correlation between MZ twins is twice as large as the correlation between DZ twins) or common environmental influences (i.e., the correlation between MZ and DZ twins is equivalent) for many regions. However, MZ and DZ twin correlations were not different from zero for some regions, indicating strong unique environmental influence. Phenotypic correlations between waves 1 and 2 ranged from .26 (right superior frontal gyrus [A8dl]) to .78 (right inferior temporal gyrus [A20cv]) and .28 (left cingulate gyrus [A23v]) to .80 (right inferior temporal gyrus [A20cv]) for ReHo and fALFF values, respectively.

### Model selection

Estimates of genetic and environmental influences on wave 1 and 2 ReHo/fALFF values were obtained from bivariate twin models specifying Cholesky decompositions (correcting for age, sex, and mean framewise displacement). Results are presented for each ROI based on the candidate model with the lowest AIC (indicative of a better fit). AE models returned the lowest AIC value for most ROIs (ReHo 171/210 ROIs; fALFF 145/210 ROIs), with CE (ReHo 34/210; fALFF 61/210) and E (ReHo 5/210; fALFF 4/210) models retained for the remaining ROIs. However, we note that AIC differences were minimal between competing AE and CE modes for many ROIs, and recommend caution when interpreting such genetic and common environment variance estimates. ROIs in which the difference in AIC values between the two best fitting models was less than two (indicating substantial support for both models (Burnham and Anderson 2002)) are signified (**STables 3-8**), and results from full ACE models are presented in **STables 9-10**.

### Raw variance components

Raw (i.e., unstandardised) variance component estimates for ReHo and fALFF values at waves 1 and 2 are presented in **STables 3-4**. Increases in total variance between the waves (denoted by non-overlapping 95% confidence intervals) were found for ReHo values of three postcentral gyrus ROIs (PoG L 4 1, PoG R 4 1, PoG L 4 4), driven by increased genetic variance. Conversely, estimates of total phenotypic variance decreased between the waves for ReHo values of 20 regions and fALFF values of 22 regions. These regions were located across the frontal, parietal and occipital lobes, with decreased genetic and environment variance amounts underlying the reduction in total phenotypic variance. Raw variance component amounts were consistent between waves 1 and 2 for the remaining ROIs.

### Standardised variance components (% of total variance)

Estimates of genetic variance standardised by total phenotypic variance (i.e., heritability; **Fig. 4-5, STables 3-4**) were of a similar magnitude between waves 1 and 2 (ReHo 17-73% and 0-66%; fALFF 17-70% and 1-66%); however, there was a greater proportion of wave 2 estimates with lower bound 95% confidence intervals spanning zero. Moderate to strong heritability estimates (>50%) were found at both waves for ReHo values of bilateral ROIs in the middle temporal, inferior temporal, and cingulate gyri. Moreover, moderate to strong heritability estimates were found at both waves for fALFF values of bilateral ROIs in the superior frontal, middle frontal, middle temporal, and inferior temporal gyri. Differences in heritability estimates between waves were observed for some ROIs (e.g., ReHo of the right inferior frontal gyrus [A44v]); however, confidence intervals for wave 1 and 2 heritability estimates were wide and overlapped between waves for all ROIs. Estimates of common environment variance were slightly stronger at wave 1 (ReHo: 20-60%; fALFF: 27-62%) than wave 2 (ReHo: 13-42%; fALFF 9-52%), with wide 95% confidence intervals for all estimates. The strongest common environment variance estimates (≈ 40-50% of total phenotypic variance at both waves) were found for ROIs in the inferior temporal (ReHo & fALFF), orbital (ReHo) and middle frontal (fALFF) gyri. Weak genetic and common environmental (and conversely strong unique environmental) influences were found for some regions, notably for ReHo values of parahippocampal gyrus ROIs (i.e., A35/36c right, TL right, TH left).

**Fig. 4.**
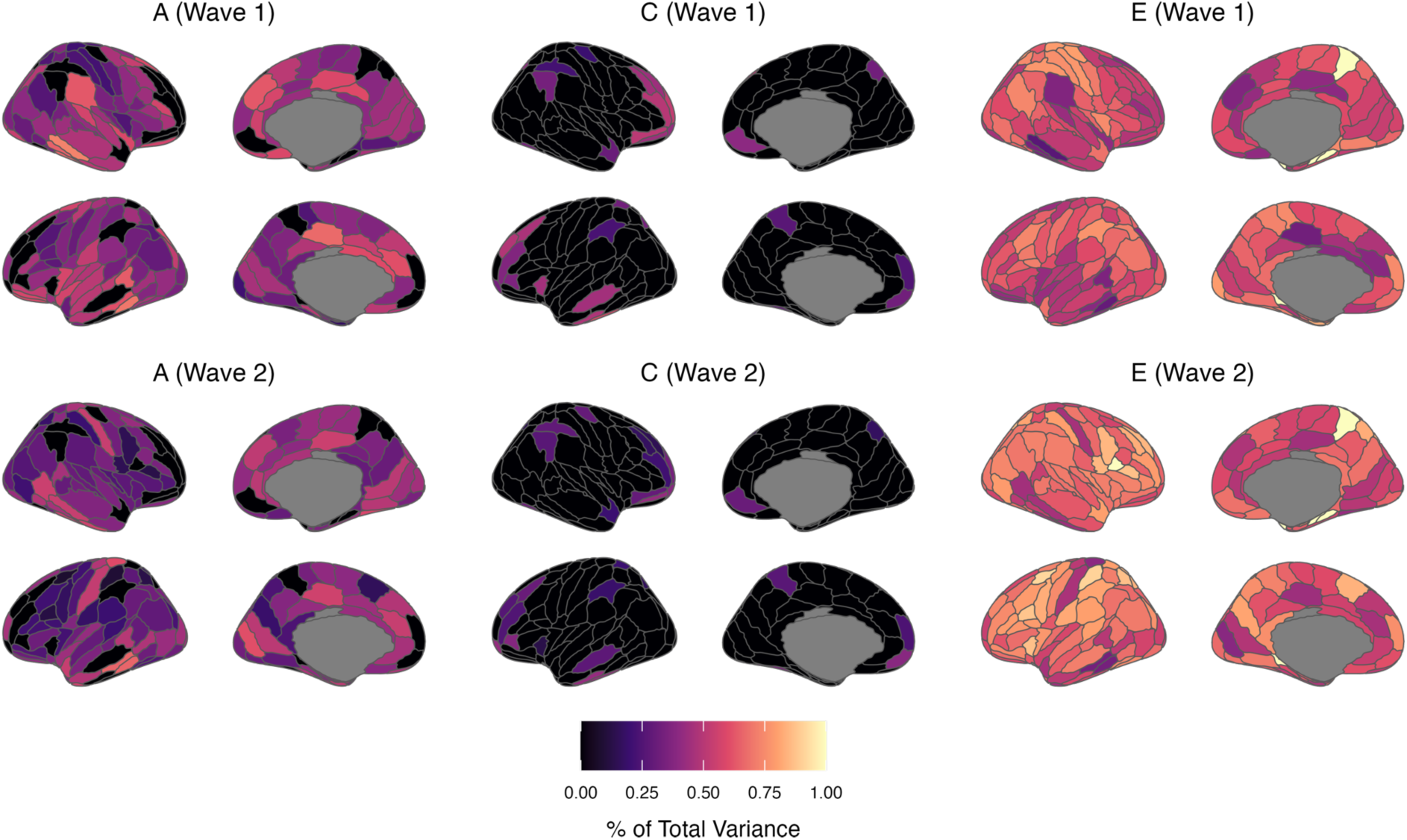
Proportions of total phenotypic variance in regional ReHo measures attributed to additive genetic (A), common/shared environment (C), and unique environment (E) latent factors (based on lowest AIC value model). Estimates are presented in **STable 3** with 95% maximum-likelihood confidence intervals.

**Fig. 5.**
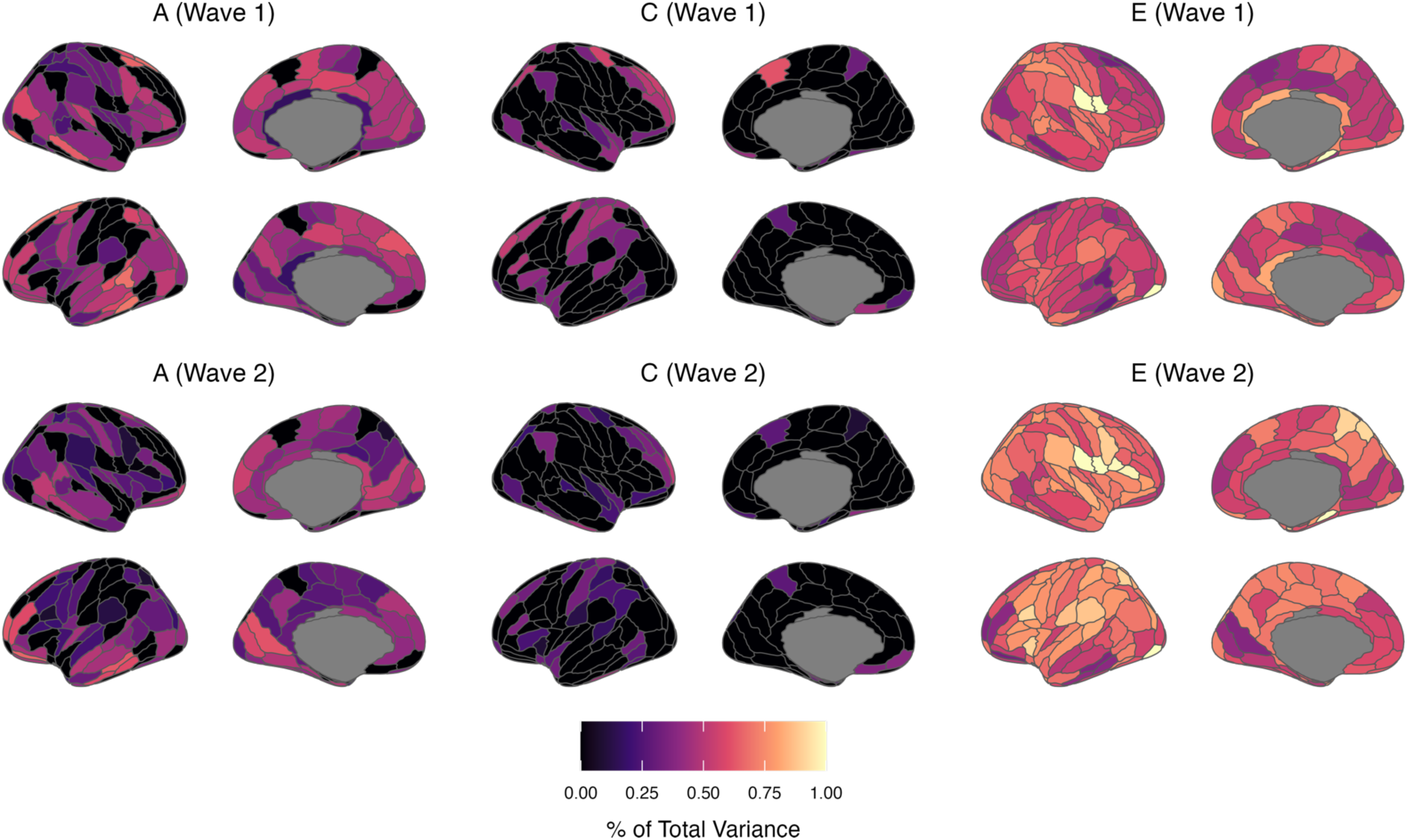
Proportions of total phenotypic variance in regional fALFF measures attributed to additive genetic (A), common/shared environment (C), and unique environment (E) latent factors (based on lowest AIC value model). Estimates are presented in **STable 4** with 95% maximum-likelihood confidence intervals.

### Continuing and new genetic and environmental influences

**Figures 6-7, STables 5-6** present parameter estimates from bivariate twin models specifying Cholesky decompositions. Estimates are standardised and squared, representing the proportion of total phenotypic variance attributed to each factor at waves 1 and 2 (summing to 1 or 100% at each wave). Results showed that new genetic influences emerged at wave 2 (i.e., path *a*_22_, **Fig. 2b**) for only a small number of ROIs (ReHo: 12 ROIs; fALFF: 25 ROIs; up to 49% of wave 2 total phenotypic variance). These ROIs were located all over the cortex, though new genetic influences on fALFF values were especially prominent in frontal and occipital lobe regions. In the remaining regions, the latent genetic factor at wave 1 (i.e., *A*1 in **Fig. 2b**) was sufficient to account for genetic variance in ReHo/fALFF values at wave 2, indicating continuing or overlapping genetic influence between waves 1 and 2. Continuing genetic influences were strongest for bilateral temporal and occipital lobe regions (ReHo and fALFF). However, 95% confidence intervals for estimates of overlapping genetic variance were wide and spanned zero for many other regions, likely due to our small sample size. We found little support for continuing genetic influences increasing in strength between the waves; however, continuing genetic influences on ReHo and fALFF values decreased in strength from waves 1 to 2 (i.e., *a*_21_ < *a*_11,_ **Fig. 2b**) in some regions. This deamplification of continuing genetic influences between the waves was present across the cortex without a clear pattern. Approximately half of the regions showing new genetic influences at wave 2 also showed a deamplification of stable genetic influences between the waves.

**Fig. 6.**
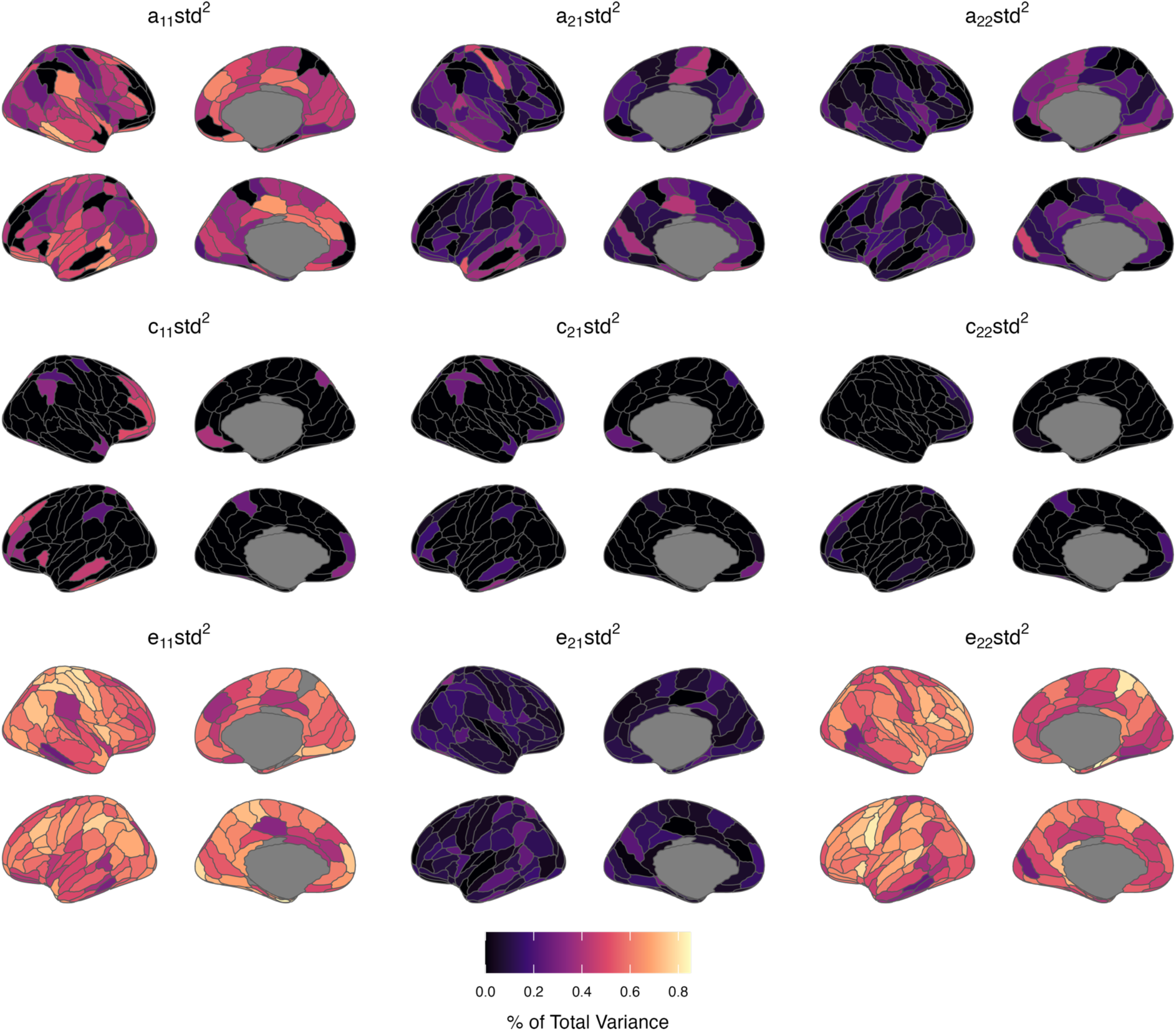
Proportions of total phenotypic variance in regional ReHo measures attributed to additive genetic (A), common/shared environment (C), and unique environment (E) latent factors (based on lowest AIC value model). Estimates reflect standardised path coefficients from bivariate twin models specifying Cholesky decompositions (squared to represent the proportion of total phenotypic variance in a trait attributed to the latent factor from which the path coefficient originates - see **Fig. 2b** for a diagram of the factors and paths). Estimates sum to 1 or 100% at each wave and are presented in **STable 4** with 95% maximum-likelihood confidence intervals.

**Fig. 7.**
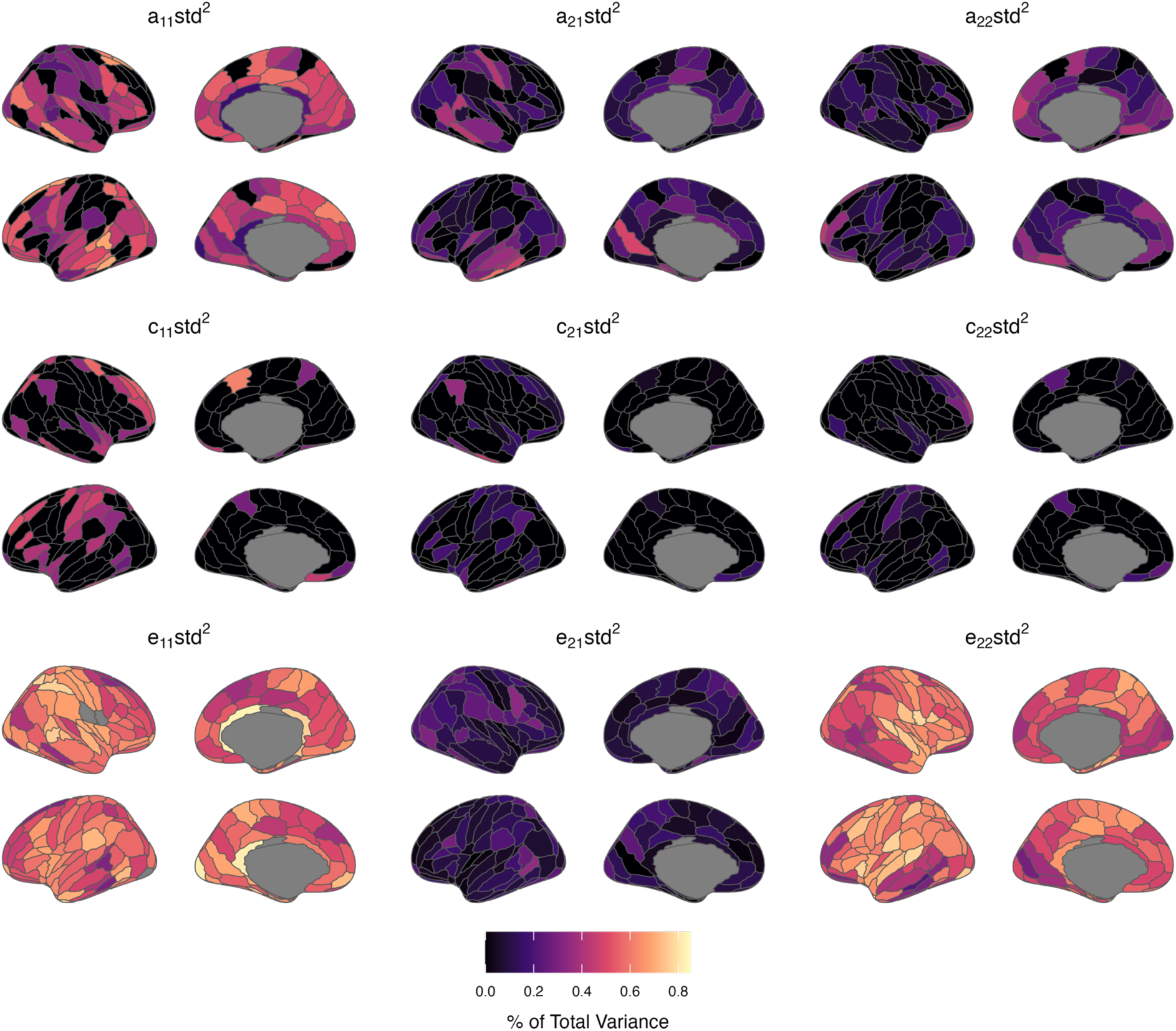
Proportions of total phenotypic variance in regional fALFF measures attributed to additive genetic (A), common/shared environment (C), and unique environment (E) latent factors (based on lowest AIC value model). Estimates reflect standardised path coefficients from bivariate twin models specifying Cholesky decompositions (squared to represent the proportion of total phenotypic variance in a trait attributed to the latent factor from which the path coefficient originates - see **Fig. 2b** for a diagram of the factors and paths). Estimates sum to 1 or 100% at each wave and are presented in **STable 5** with 95% maximum-likelihood confidence intervals.

Regions showing new common environmental variance at wave 2 (i.e., path *c*_22_, **Fig. 2b**) were sparse and most prominent for fALFF values of the right middle frontal gyrus regions (A9/46d, A46, A9/46v). Consequently, common environment factors influencing ReHo/fALFF were stable between waves 1 and 2 for most ROIs. These effects were strongest for ReHo and fALFF values of lateral frontal, temporal, and parietal regions. Similar to results for genetic parameter estimates, we found evidence of stable common environmental decreasing (but not increasing) in strength between the waves. ROIs with decreasing stable common environmental influences (ReHo: 10 ROIs; fALFF: 24 ROIs) were found predominantly in frontal (ReHo, fALFF) and parietal (fALFF) regions. Estimates of stable unique environmental influences between the waves were slightly weaker than corresponding genetic or common environmental influences. However, substantial estimates of novel unique environmental influence (i.e., *e*_22_, which includes measurement error) were found for ReHo and fALFF values, ranging from 25-82% and 23-80% of total wave 2 phenotypic variance, respectively.

### Associations between ReHo/fALFF and mental health and cognition measures

Linear mixed-effects models using data from waves 1 and 2 found small and positive associations between anxiety/depressive symptoms (i.e., SCAS, SPHERE-21, & SMFQ measures) and ReHo or fALFF values of several temporal lobe ROIs (**STables 7-8**). Notably, fALFF of the left fusiform gyrus (A20rv) and left inferior temporal gyrus (A20r) ROIs was positively associated with all three anxiety and depression measures. Further, ReHo values of the left fusiform gyrus (A20rv) ROI were positively associated with SPHERE-21 and SCAS score, and ReHo values of the left parahippocampal gyrus (A35/36) ROI were positively associated with SPHERE-21 and SMFQ score. No associations between regional ReHo or fALFF values and Crystalized Cognition Composite score remained following multiple testing correction.

## Discussion

To better understand differences in brain function among adolescents, we studied the influence of genetic and environmental factors on resting-state fMRI measures of brain function (specifically ReHo and fALFF) in a group of adolescent twins over two occasions. Results showed that genetic and environmental factors influenced brain function in almost all 210 cortical regions examined. Moreover, genetic and common environmental factors influencing ReHo and fALFF values at wave 1 (9-14 years) also influenced values at wave 2 (10-16 years) for many regions. However, influences of genetic and common environmental factors varied across the cortex, exhibiting different patterns in different regions. Furthermore, we found new (i.e., independent) genetic and environmental influences on brain function at wave 2, again with regional patterns. Additionally, we found weak associations between anxiety and depressive symptoms and local brain function in several regions of the ventral temporal lobe.

We found a wide range of heritability estimates for ReHo and fALFF values across the cortex (ReHo 17-73% [wave 1] and 0-66% [wave 2]; fALFF 17-70% [wave 1] and 1-66% [wave 2]). This is consistent with the estimate for whole-brain ReHo heritability (67%) reported by Adhikari et al., (2020) and in line with estimates from functional connectivity studies (Glahn et al. 2010; Korgaonkar et al. 2014; Adhikari et al. 2018; Teeuw, Brouwer, Guimaraes, et al. 2019; Reineberg et al. 2020). High heritability estimates (60-80%) have been reported for functional connectivity measures (Fornito et al. 2011; Fu et al. 2015; Achterberg et al. 2018); however, these estimates may be overestimated due to sampling error associated with small sample size (i.e., 30-60 pairs). Heritability estimates for ReHo and fALFF measures are lower than those for structural MRI measures (Lenroot et al. 2009; Jansen et al. 2015; Maes et al. 2023). The greater unreliability of fMRI measures compared to structural MRI measures (Buimer et al. 2020) likely underlies, at least partly, the lower heritability estimates of functional measures.

In a sample of adolescent twins scanned at ages 13 and 18, Teeuw et al. (2019) found stronger genetic influences on functional connectivity within the visual, frontoparietal, and salience networks compared to the language, dorsal attention, and sensorimotor networks, with common environment influences on default mode network connectivity. Although heritability estimates for ReHo/fALFF and functional connectivity are not strictly comparable, we find some similarities with our results. Indeed, estimates of genetic influence on ReHo and fALFF values were strongest for ROIs located in the frontoparietal network (ROI-network correspondence based on the mapping of Brainnetome ROIs to the 7 network parcellation of Yeo et al., (2011), provided online at https://atlas.brainnetome.org/). Further, heritability estimates were weakest for ROIs in the somatomotor, dorsal attention, ventral attention networks (ReHo), and limbic networks (fALFF). In contrast to the common environmental influences Teeuw et al. (2019) reported, we found relatively strong genetic influences on Reho and fALFF values for select default mode network ROIs. We further found generally low to moderate heritability estimates for ROIs in the visual network, for which Teeuw et al. (2019) found strong genetic influences.

We found stable genetic or common environmental influences on ReHo/fALFF between waves 1 and 2 for most regions, similar to that found by Teeuw et al., (2019) for functional connectivity estimates. However, we found a greater incidence of new (i.e., independent) genetic influences at our second time point than Teeuw et al. (2019), potentially due to our larger sample. The emergence of novel genetic influences across adolescence may reflect targeted genetic influences on specific brain regions related to the behavioural and cognitive changes of adolescence. Such “areal specialisation” of the cortex was posited to explain the emergence of novel genetic influences on cortical thickness during adolescence (Teeuw, Brouwer, Koenis, et al. 2019). Indeed, estimates of new genetic influence at wave 2 were greatest for fALFF values of frontal lobe regions, with estimates for ReHo values greatest for fusiform gyrus regions (amongst others). However, we must note that regions not typically associated with behavioural/cognitive change (predominantly occipital lobe ROIs) also showed novel genetic influences on ReHo and fALFF values at wave 2.

We found stable common environmental factors to influence ReHo/fALFF of several ROIs, with little support for new common environmental factors at wave 2. Shared environmental variance likely reflects the influence of various environmental factors, including family environment and education system. There is some evidence of shared environmental influence on functional connectivity in adolescents and young adults (Yang et al. 2016; Teeuw, Brouwer, Guimaraes, et al. 2019), but little evidence exists for adults (Sinclair et al. 2015; Elliott et al. 2019; Reineberg et al. 2020). However, results from both the current and past examinations are limited by sample size. Indeed, the shared environment influences reported may represent a lack of statistical power to distinguish between genetic and shared environmental effects (reflected in the minimal AIC differences between AE and CE models and low twin correlations for many of these connections). Consequently, larger sample sizes are required to confirm shared/common environmental influences on functional connectivity.

For some ROIs, there was a lack of genetic or common environment variance. This predominant influence of unique environmental factors may reflect regional brain function especially sensitive to non-genetic influences (i.e., stochastic biological effects or idiosyncratic experiences, such as school stress and physical activity). However, rs-fMRI measures are inherently noisy, and as estimates of unique environmental influence include measurement error, such environmental variance may reflect measurement unreliability. This is commonly assessed using test-retest reliability estimates (Elliott et al. 2019; Ge et al. 2024), though the absence of a test-retest sample in the QTAB dataset precludes this. Interestingly, ROIs in which variance was attributable entirely to unique environmental factors (ReHo: 5 ROIs; fALFF: 4 ROIs) did not show especially low phenotypic correlations between wave 1 and 2 values (*r*_ph_ .40-.54; a proxy for long-term reliability), suggesting that measurement unreliability is not exclusively responsible for the strong unique environmental variance found. This is further supported by evidence showing that ReHo and fALFF metrics show greater reproducibility than functional connectivity measures (Golestani et al. 2017; Cahart et al. 2023).

ReHo and fALFF displayed region-specific differences between males and females, with sex differences slightly more pronounced at wave 2. Previous studies have reported sex differences in rs-fMRI measures in children (Alarcón et al. 2015; Teeuw, Brouwer, Guimaraes, et al. 2019; Yoon et al. 2021; Liang et al. 2022) and adults (Biswal et al. 2010; Zhang et al. 2016; Ritchie et al. 2018). Given the myriad methodological differences between these studies, it is challenging to elucidate consistent sex effects on brain activity and function measures. However, an overall conclusion may be that any actual sex effects are likely extremely subtle (Eliot et al. 2021). We did not find widespread effects of age on ReHo or fALFF, likely due to our narrow participant age range. We found small associations between mean framewise displacement (i.e., head motion) and ReHo/fALFF values of many regions, even though participants with mean framewise displacement greater than .3mm were excluded. These associations reflect the pervasiveness of head motion in rs-fMRI measures and highlight the importance of reporting and controlling for motion.

Exploratory analyses between ReHo/fALFF measures and self-report scores of anxiety and depressive symptoms highlighted several regions of the ventral temporal lobe. While studies examining neural substrates of depression and anxiety typically focus on emotional (dys)regulation, recent research highlights the role of conceptual processing and semantic cognition in developing depressive and anxiety disorders (González-García & Visser, 2023). The ventral temporal lobe underlies conceptual processing/semantic memory (Ralph et al. 2017), enabling understanding verbal and nonverbal conceptual information and modulating higher-level social behaviour through connections to other limbic structures. Past examinations of ReHo/fALFF and behavioural associations, typically case-control studies in small samples (*n* < 100), have implicated a range of cortical regions (Oathes et al. 2015; Shen et al. 2020), which have included the temporal lobe (Zhao et al. 2023). Further, associations between depression-anxiety scores and occipital-temporal cortical surface area have been reported in young adults (Couvy-Duchesne et al. 2018). As the sample size required to detect reproducible brain-behaviour phenotypic associations is many times larger (Marek et al. 2022), studies in larger samples are required to extend the results of such an exploratory analysis.

The results of the present study should be considered in the context of several limitations. Firstly, as indicated by the wide confidence intervals of all estimates, our small sample size reduces the precision of the presented genetic and environmental variance estimates. Secondly, selecting a best-fitting model (i.e., ACE, AE, CE, or E) is a complex and somewhat nuanced task which does not conform to easily applied criteria. Our practice of selecting the candidate model with the lowest AIC value is a simplistic approach that facilitates examining many variables using a single criterion. Further, twin models using Cholesky decomposition show discrepancies between numerical and theoretical Type I error rates and slightly biased parameter estimates (Verhulst et al. 2019). An alternative approach is to estimate variance components directly; however, this approach can return nonsensical estimates (e.g., negative variance estimates) due to sampling variability in small studies. Hence, to aid the interpretability of our longitudinal dataset, we elected to retain the Cholesky decomposition approach. Lastly, the genetic and environmental variance estimates for ReHo and fALFF values presented here are likely dependent (to some degree) on the preceding methodological steps (e.g., rs-fMRI preprocessing, selection of cortical parcellation).

Results from the current study highlight an intricate pattern of genetic and environmental influences underlying variation in ReHo and fALFF measures of brain function during early-to-mid adolescence (9-16 years). These effects are largely stable between the two time points examined; however, changes in the genetic and non-genetic factors influencing ReHo and fALFF were found between the waves for some regions. Exploratory analyses suggest a link between these measures of brain function and mental health during adolescence. Well-powered longitudinal studies are needed to understand these variance patterns further and their implications regarding adolescent brain development.

## Acknowledgements

The Queensland Twin Adolescent Brain (QTAB) study was supported by the National Health and Medical Research Council (NHMRC; grant number 1078756), as was SEM (grant numbers APP1172917 and APP2025674). Special thanks to Margie Wright for her leadership of the QTAB project. Thanks also to Narelle Hansell and Paul Thompson for their involvement in the QTAB project. Most of all, we are forever grateful to the twins and their families for their willingness to participate in our studies.

